# Distributed representation of context by intrinsic subnetworks in prefrontal cortex

**DOI:** 10.1101/074880

**Authors:** Michael L. Waskom, Anthony D. Wagner

## Abstract

Human prefrontal cortex supports flexible decision-making by representing abstract information about task context. The organizational basis of these context representations, and of representations underlying other higher-order processes, is unknown. Here, we use multivariate decoding and analyses of spontaneous correlations to show that context representations are distributed across subnetworks within prefrontal cortex. Examining targeted prefrontal regions, we found that pairs of voxels with similar context preferences exhibited spontaneous correlations that were approximately twice as large as those between pairs with opposite context preferences. This subnetwork organization was stable across task-engaged and resting states, suggesting that abstract context representations are constrained by an intrinsic functional architecture. These results reveal a principle of fine-scaled functional organization in association cortex.

## Introduction

The cerebral cortex exhibits functional organization at multiple spatial scales. At a coarse scale, the cortex is parcellated into functional areas (1) that coordinate as networks through long-range connections (2–4). These regions represent and compute information with functional circuitry that is organized more finely. Some fine-scaled principles have been described in detail, particularly for sensory cortex (5, 6). In contrast, the subregional organization of prefrontal cortex (PFC), which gives rise to higher-order processes including attention, decisionmaking, and goal-directed behavior (7, 8), remains largely uncharacterized. In particular, it is unknown whether PFC representations are encoded within an equipotential system or constrained by an intrinsic functional architecture.

Sensory cortex can be mapped by parametrically varying stimulus attributes and measuring changes in the neural response, but complex and dynamic response properties limit the effectiveness of this approach for mapping PFC and other association regions. An alternate strategy is to leverage spontaneous variability in neural activity. Neural responses to repeated presentations of sensory stimuli, or in the absence of stimulation and explicit task demands, exhibit variability that is attributed to ongoing spontaneous activity (9–12). Traditional analyses consider spontaneous activity to be noise, but there is increasing evidence that shared spontaneous variability is a signature of functional organization (13). Analyses of spontaneous correlations have been used to identify multiple large-scale functional networks (3, 4, 14) and boundaries between functional areas (15, 16) in human and nonhuman primate association cortex. The spontaneous correlation structure also mirrors established fine-scaled principles of organization in visual cortex, such as preferences for retinotopic position (17) or stimulus orientation (18). We therefore leveraged spontaneous activity to examine fine-scaled functional organization in human PFC.

We focused our investigation on prefrontal representations that enable flexible decision-making. Across two experiments, we scanned subjects with high-resolution fMRI while they performed tasks that demanded selective integration of noisy sensory evidence according to a shifting decision rule, or context. During rule-based decisionmaking, distributed patterns of activation in lateral prefrontal cortex encode a representation of the context (19–21). We have previously reported that context representations can be localized to the inferior frontal sulcus (IFS), a prefrontal region defined through large-scale analyses of spontaneous correlations (22, 23). Here, we use multivariate decoding and analyses of subregional spontaneous correlations to show that context representations are organized across intrinsic subnetworks within the IFS. We propose that functionally specific subnetworks could be a general feature of cortical organization and that they may underlie distributed representations in other cognitive domains.

## Results

In Experiment 1, subjects viewed a bivalent random dot stimulus and were cued to discriminate either the direction of coherent motion or the more prevalent color (Supplemental Figure 1A). We first sought to identify distributed patterns representing the task context (i.e., the motion vs. color rule). Guided by our previous findings (22), we applied linear classifiers to patterns of activation within the bilateral IFS (Figure 1A). Decoding performance exceeded chance at both the group (*t*_13_ = 7.66, *P* < 0.001; Figure 1B) and subject (11/14 subjects *P* < 0.05, permutation test) levels. We also evaluated a region in medial PFC (MFC; Figure 1A) that is widely implicated in cognitive control processes (24) but that we did not expect to strongly represent task context. Decoding performance in MFC did not exceed chance (*t*_13_ = 1.11, *P* = 0.29; 1/14 subjects *P* < 0.05, permutation test; IFS vs. MFC paired *t*_13_ = 7.10, *P* < 0.001; Figure 1B), which confirmed the relative importance of the IFS for context representation.

**Figure 1.**
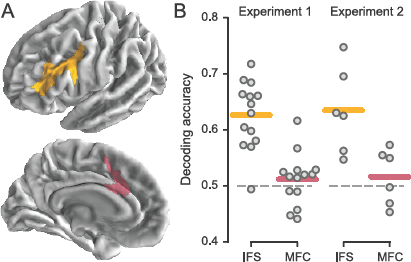
Decoding task context in targeted PFC regions. (A) Region labels as defined on the group surface. Labels were reverse-normalized to individual subject brains before analysis. (B) Cross-validated accuracy for decoding task context (motion vs. color rule in Experiment 1; orientation vs. color rule in Experiment 2). Each point corresponds to the accuracy for an individual subject; horizontal lines indicate group means.

To evaluate the functional organization of these representations, we inverted the decoding models (see Methods) to obtain voxelwise estimates of context preference. When plotted on the cortical surface, voxels with similar preferences appeared organized into relatively fine-scaled, but not random, clusters (Figure 2 and Supplemental Figure 2). To quantify the spatial organization, we asked whether it was possible to reliably predict a voxel’s preference from those of its neighbors at different distances (Supplemental Figure 3). This analysis suggested that voxel preferences were both clustered and interdigitated relative to a simulated random organization, with an estimated upper bound for the local neighborhood size of 15.7 ± 2.5 mm (mean ± s.d.).

**Figure 2.**
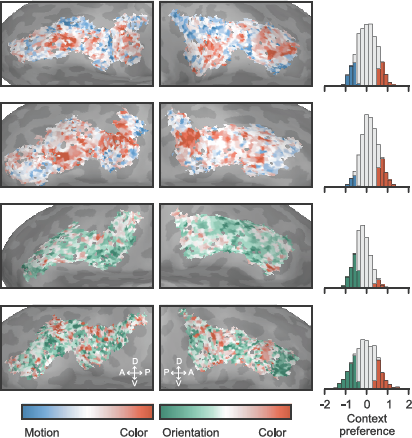
Context preferences in the IFS. Results are shown for four example subjects (for all subjects, see Supplemental Figures 2 and 4). Left and right columns correspond to left and right hemispheres, respectively. Colored histogram bars indicate voxels selected, using a permutation test, for analysis of spontaneous correlations.

Our main question was whether patterns of spontaneous activity reflect the functional organization of distributed context representations within IFS. To answer this question, we first used a permutation test to identify voxels with relatively strong preferences for color or motion trials (Figure 2 and Supplemental Figure 2). For each selected voxel, we then regressed out a model of task variables to produce an estimate of spontaneous activity and computed the timeseries correlation between each voxel pair (11, 25). Using multidimensional scaling (MDS) to reduce the dimensionality of the spontaneous correlation structure, we observed clustering of voxels with similar context preferences (Figure 3A and Supplemental Figure 5). This clustering implies that voxels with similar context preferences exhibit relatively elevated spontaneous correlations. To formally test this relationship, we computed the mean timeseries correlations between all pairs of voxels with the same binary context preference and all pairs with different context preferences. Same-context spontaneous correlations were approximately twice as large as different-context spontaneous correlations (paired *t*_13_ = 9.04, *P* < 0.001; 14/14 subjects *P* < 0.05, permutation test; Figure 3B). Stronger spontaneous correlations between voxels with similar context preferences indicates that they are organized into functionally specific subnetworks.

**Figure 3.**
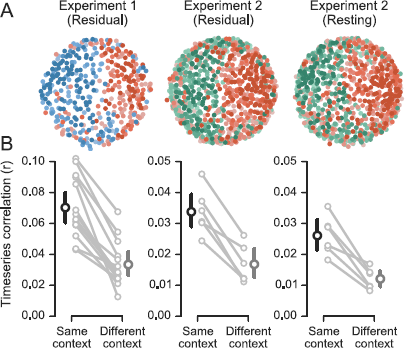
Spontaneous correlations reveal subnetworks that represent task context. (A) The relationship between spontaneous correlation structure and voxel-wise context preferences are shown for two example subjects. The positions of voxels were determined using a multidimensional scaling of the spontaneous correlation matrix; the voxel context preferences are shown with the colormaps from Figure 2. The same subject is shown for both Experiment 2 analyses. (B) Mean same-context and different-context spontaneous correlations. Large points and error bars show the group means and 95% confidence intervals; correlation values for individual subjects are connected with a line.

In Experiment 2, we sought to determine whether these subnetworks are apparent when spontaneous activity is measured without sensory stimulation or explicit task demands. We therefore scanned an additional set of subjects both in the resting-state and during performance of a context-dependent perceptual discrimination task. In this task, subjects were cued to determine either the more prevalent orientation or color of a field of small “sticks” (Supplemental Figure 1B). As in Experiment 1, we applied linear classifiers to identify a distributed representation of task context in the IFS (*t*_5_ = 4.10, *P* = 0.009; 4/6 subjects *P* < 0.05, permutation test; Figure 1B), which was not observed in the MFC (*t*_5_ = .86, *P* = 0.43; 2/6 subjects *P* < 0.05, permutation test; IFS vs. MFC paired *t*_5_ = 3.96, *P* = .01; Figure 1B). The spatial organization of context preferences in Experiment 2 also provided evidence for fine-scaled clustering and interdigitation (Figure 2 and Supplementary Figures 3 and 4).

The within-IFS spontaneous correlation structure was highly stable between task-engaged and resting states (Supplemental Figure 6). To formally test the stability of the correlations across task and rest, we evaluated the similarity of voxelwise correlation matrices derived from the two kinds of measurements. This test confirmed the similarity of the correlation structures (mean *r* = 0.80, *t*_5_ = 35.93, *P* < 0.001; 6/6 subjects *P* < 0.05, Mantel test). Moreover, we did not observe any general trends for elevated correlations in one of the two states. Although we cannot rule out context-dependent changes in subnetwork structure, we conclude that task-engaged and resting-state measurements of spontaneous correlations provide convergent information.

The results of Experiment 2 replicated the finding that distributed representations of context are expressed across fine-scaled subnetworks in IFS (Figure 3). As in Experiment 1, spontaneous correlations identified from the residual signal in a task-engaged state were approximately twice as large for voxel pairs with the same context preference relative to voxel pairs with different context preferences (paired *t*_5_ = 7.43, *P* < 0.001; 6/6 subjects *P* < 0.05, permutation test; Figure 3B). Importantly, this was also the case when spontaneous correlations were identified in resting-state scans (paired *t*_5_ = 5.96, *P* = 0.002; 6/6 subjects *P* < 0.05, permutation test; Figure 3B). Qualitative features of the subnetwork organization, as apparent in the MDS plots (Figure 3A and Supplemental Figure 5), were also similar between task-engaged and resting states.

The identification of fine-scaled subnetworks in the resting-state suggests that they are an intrinsic feature of prefrontal organization. Because the resting-state acquisitions alternated with task scans, though, it is possible that subnetworks dynamically emerge during task performance and persist for several minutes following the offset of a task. This alternative account would predict that subnetworks should not be identifiable in the first resting-state scan of each session, which was acquired before task performance, and potentially that subnetwork strength should increase over time. To evaluate these predictions, we recomputed the difference in same-context and different-context correlations for each of the four resting-state acquisition times (Supplemental Figure 7). This analysis confirmed that subnetworks were identifiable in the first resting-state scan (6/6 subjects *P* < 0.05, permutation test) and provided no consistent evidence for change in subnetwork strength over time.

We also evaluated whether the relationship between context preference and spontaneous correlation could be fully attributed to short-range correlations between spatially clustered voxels with similar preferences. Specifically, we recomputed the mean same-context and different-context correlations while excluding voxel pairs below different thresholds of distance along the cortical surface (Supplemental Figure 8). The results indicated that subnetworks extend over relatively long distances, as the relationship remained significant even when excluding voxel pairs situated closer than 40mm (13/14 and 6/6 subjects *P* < 0.05, permutation test). This analysis implies that our findings cannot be accounted for by spatial autocorrelation in the BOLD signal.

## Discussion

Our results reveal an intrinsic organizational basis for the distributed representation of task context, a higher-order process that underlies the capacity for flexible decision-making (7). Our main result is that voxels with similar context preferences exhibited spontaneous correlations that were approximately twice as large as the those between voxels with opposite preferences. This provides evidence for the existence of functionally specific subnetworks within lateral prefrontal nodes of the large-scale frontoparietal control network. Like large-scale functional networks, subnetworks can be identified based on the spontaneous correlation structure measured in resting-state activity or in residual activity during task performace. Subnetwork functional organization is fine-scaled, relative to the encompassing region, and the spatial organization appears to be idiosyncratic across subjects.

Subnetwork organization qualitatively resembles densely interconnected “stripes” of neurons that have been observed using anatomical tracers in nonhuman primate PFC (26). Computational models of working memory and cognitive control suggest that these stripes are involved in organizing prefrontal representations (27), but their functional relevance has not been empirically determined. In line with model predictions, our results suggest that these design principles are relevant for structuring abstract representations in human PFC. Despite this qualitative agreement, our spatial clustering analyses indicate that context preferences are shared across larger spatial extents than can be accounted for by the stripe sizes previously observed in nonhuman primates (26). This may represent a difference between species or the effect of blurring within our voxels. Quantitative between-species comparisons will require functional imaging data with a higher spatial resolution.

The relationship between context representation and spontaneous correlations suggests a relatively small number of subnetworks in the IFS. If the activity of each subnetwork provides information about one context, this implies limits on the capacity of context representation that may seem incongruent with the broad diversity of tasks that humans are capable of performing. However, humans exhibit striking limits in their attention and working memory capacity (28), and these constraints are broadly related to general cognitive abilities (29). Humans’ abilities to actively represent and evaluate rules appears similarly limited (30). The functional properties of prefrontal subnetworks may provide insight into the neural mechanisms that underlie constraints on cognitive processing capacity.

Multivariate decoding approaches are increasingly used to test theories about higher-order representations (31), but the organizational principles that underlie successful decoding in association cortex have remained unclear. Our data indicate that, for context decoding in lateral PFC, information is expressed across patches on the order of tens of millimeters. Moreover, the elevated short-and long-range spontaneous correlations within and between these patches suggest that they comprise functional units. It is an open question whether this organization is specific to the representation of context in lateral PFC or if it is a general pattern in association cortex. We note that large-scale networks exhibit consistent organization across the brain (3, 4), and we suggest that fine-scaled subnetworks may likewise exist in other regions and could similarly provide structure for other high-level cognitive representations.

Analyses of spontaneous correlations identify highly reproducible large-scale networks in the human brain (3, 4), but the relevance of these measurements to finer scales of organization has remained controversial. Some aspects of fine-scaled organization in sensory cortex are reflected in the spontaneous correlation structure, including retinotopic eccentricity biases (17) and orientation maps (18). Analyses that identify discontinuities in spontaneous correlation structure have also been applied in PFC to characterize boundaries in subregional organization (15, 16). Our results extend these previous findings in two important ways. First, they show that the fine-scaled spontaneous correlation structure in PFC constrains task-directed representations of abstract context information. Second, they indicate that subregional organization is not characterized just by discrete areas but by a subnetwork structure. These findings demonstrate the effectiveness of spontaneous correlation analyses as a tool for studying the fine-scaled functional organization of association cortex and emphasize the importance of functionally specific network organization in the human brain at multiple spatial scales.

## Acknowledgments

This work was supported by the Stanford Center for Cognitive and Neurobiological Imaging and the Knut and Alice Wallenberg Foundation (Network Initiative on Culture, Brain, and Learning). M.L.W. was supported as an NSF IGERT Traineeship Recipient under Award 0801700. We thank Serra Favila and Karen LaRocque for assistance with data collection and helpful conversations, Bob Dougherty for assistance with the high-resolution pulse sequence, and Roozbeh Kiani, Brett Foster, and Thomas Yeo for comments on an earlier version of the manuscript.

## Methods

### Subjects

Subjects were healthy right-handed members of the Stanford community and gave informed written consent in accord with the Stanford University Institutional Review Board. All subjects had normal or corrected-to-normal visual acuity and normal color vision. Subjects received $10/hour for behavioral sessions and $20/hour for scanning, and they additionally received a monetary bonus based on their performance during the imaging sessions.

In Experiment 1, 20 subjects were recruited. Of this group, one was excluded early in the training session for poor color vision, one was excluded after the training session for performing at chance, and three ended the scanning session early due to fatigue or illness. One additional subject was excluded after initial data analysis suggested experimenter error during data acquisition; the decision to exclude this subject was made before performing these analyses. The analyses reported here reflect data from the 14 remaining subjects (19-29 years old; 7 females). Each subject participated in one behavioral training session and one imaging session for approximately 3.5 total hours participation. In Experiment 2, eight subjects were recruited; all had extensive experience being scanned, one subject was author MLW, and one subject had participated in Experiment 1. Of this group, one subject was excluded for early exit from the scanning session, and one subject was excluded due to data loss during image reconstruction. The analyses reported here reflect data from the remaining six subjects (24-29 years old; 3 females). Each subject participated in two behavioral training sessions and two imaging sessions for approximately 5 total hours participation.

### Stimuli and Experimental Design

#### Experiment 1

The stimuli and experimental design for Experiment 1 have been previously reported in detail (23). Briefly, subjects viewed a bivalent random dot stimulus and were cued to make either a motion or a color discrimination on each trial (Supplemental Figure 1A). Motion information was controlled by displacing a selected proportion of the dots coherently either up or down on each screen refresh, while the remainder of the dots were redrawn at a random location. Color information was controlled by drawing a selected proportion of the dots in either red or green on each screen refresh. The difficulties of the color and motion discriminations were set independently for each subject using a staircase procedure in a separate training session prior to scanning. On each trial, the relevant dimension (the context) was cued by the pattern of the frame surrounding the stimulus. Two distinct patterns cued a motion trial, and two distinct patterns cued a color trial. The cue appeared 1 s before the stimulus on  of trials, it appeared concurrent with stimulus onset on  of trials, and  of trials were “cue-only” trials where the frame was presented for 1 s but was not followed by a stimulus. On trials with a stimulus, it was shown for 2 s. Subjects were instructed to respond as soon as they had made a decision and indicated their response with a button box held in the right hand. No feedback was presented during scanning. The experiment was structured into epochs with different proportions of motion and color trials (see ref (23) for details). In total, each subject performed 900 trials (600 with a stimulus), evenly divided between motion and color contexts, across 12 scanner runs.

#### Experiment 2

The stimuli and experimental design for Experiment 2 are reported in detail in the Supplemental Methods. Briefly, subjects performed a context-dependent perceptual decision-making task that was similar to the task in Experiment 1. Subjects viewed a bivalent stimulus that comprised a field of small sticks and were cued to make either an orientation or a color discrimination on each trial (Supplemental Figure 1B). Orientation and color information was manipulated by controlling the proportion of sticks drawn at either 45° or 135° and in either red or green. The difficulties of the orientation and color discriminations were set independently for each subject using performance in two separate training sessions prior to scanning. In contrast to Experiment 1, two different difficulty levels were used for each dimension during scanning. On each trial, the relevant dimension (the context) was cued by the shape of a polygon drawn at fixation. Two distinct polygons cued an orientation trial and two distinct polygons cued a color trial. In contrast to Experiment 1, the cue appeared concurrently with stimulus onset on every trial, the stimulus disappeared when subjects made their first button press response, and feedback was provided by blinking the fixation point after error trials. The relevant dimension on each trial was chosen randomly from a balanced distribution. In total, each subject performed 768 trials, evenly divided between orientation and color contexts, across 12 scanner runs in two separate scanning sessions.

#### Resting-state Scans

We collected eight separate resting-state scans from each subject. Each resting-state scan had a duration of 367.2 s. Both scanning sessions began with a resting-state scan, and the remainder were interleaved with the task runs. During resting-state scans, a black fixation cross was displayed on a grey background drawn at 30% luminance. Subjects were instructed to fixate on the cross and to let their minds wander. Fixation and wakefulness were monitored with an eye-tracking camera.

#### fMRI Acquisition

Brain imaging was performed on a 3T GE Discovery MR750 system (GE Medical Systems) using a 32-channel transmit-receive head coil (Nova Medical). In Experiment 1, partial-brain functional images were obtained using a T2*-weighted two-dimensional echo planar imaging sequence with a relatively high spatial resolution (TR = 2 s, TE = 30 ms, flip angle = 77°, 33 slices, 96 × 96 matrix, 2 × 2 × 2.3 mm voxels, axial oblique interleaved acquisition). Slices were acquired aligned with and dorsal to the superior temporal sulcus to provide coverage of frontal and parietal cortex. In Experiment 2, whole-brain functional images were obtained with high spatial and temporal resolution using a T2*-weighted multi-band 2D echo planar imaging sequence(32) (TR = 720 ms, TE = 30 ms, flip angle = 53°, 64 slices, 110 × 110 matrix, 2 mm isotropic voxels, multi-band factor = 8, axial oblique interleaved acquisition). Aside from scan duration, acquisition parameters were identical for task and resting runs. In both experiments, whole-brain high-resolution (0.9 mm isotropic voxels) Ti-weighted SPGR volumes were acquired for cortical surface modeling and across-run alignment.

#### fMRI Preprocessing

Unless otherwise specified, data from Experiments 1 and 2 were processed with the same workflow. After image reconstruction, images from Experiment 2 underwent susceptibility distortion correction using two single-band images with reversed phase-encode blips (33). This step was omitted for Experiment 1 because we did not collect fieldmap images. Next, each timeseries was realigned to its middle volume using normalized correlation optimization and cubic spline interpolation. The timeseries data for Experiment 1 were then temporally resampled using sinc interpolation to correct for slice timing differences. This step was not performed for Experiment 2 data because of the high temporal resolution and multiband acquisition. Each timeseries was then scaled with a single multiplicative factor so that the median value across time and space (within a brain mask derived from the anatomical volume) equaled 10000. Finally, the timeseries data were high-pass temporally filtered by fitting and removing gaussian-weighted running lines with an effective cycle cutoff of 128 s. Functional data were not spatially smoothed.

Separately, the T_1_-weighted anatomical volume (or average volume in the case of multiple acquisitions) was processed using Freesurfer to segment the grey-white matter boundary and construct tessellated meshes representing the cortical surface (34). A linear transformation (6 degrees of freedom) between the native functional space and the native anatomical space was estimated for each run using the mean functional image from each timeseries. This registration was estimated by optimizing a boundary-based cost function, which aligns the data with the model of grey-white boundary (35).

Timeseries data were then denoised by fitting and removing models with head motion and anatomically derived confound information. Specifically, this model included 6 motion regressors (3 rotations and 3 translations), 6 regressors representing principal components of signals extracted from the deep white matter, and indicator vectors identifying scans with signal artifacts. Anatomical confound extraction was performed by transforming the Freesurfer-segmented white matter mask into the functional space, eroding it by 3 voxels, and submitting the resulting time × voxel matrix to principal components analysis. Signal artifacts were automatically determined using the following criteria: any frame in which the motion parameters indicated a displacement relative to the previous frame larger than a threshold (0.5 mm in Experiment 1 and 0.25 mm in Experiment 2); any frame in which the median signal across the whole-brain mask deviated from the run median by more than 5 median absolute deviations (MAD); any frame in which any median slice intensity, after subtracting the whole-brain median, deviated from the run median by greater than 10 MAD. For the resting-state data, we additionally included a regressor representing the mean signal at each timepoint across a whole-brain mask determined using the Freesurfer segmentation. Whole-brain regression was omitted for the task data. These models were fit using linear regression, and the residual data were used in subsequent analyses.

The middle volume of the first functional task run served as the common space for each subject. Other frames from the first run were realigned to this space during motion correction. Frames from other runs were first realigned to that run’s middle volume and then transformed into the common space by concatenating the transformations from that run to the anatomy and the inverse of the transformation from the first run to the anatomy. Images were resampled with cubic spline interpolation. All analyses were performed in a native, voxel-based space; data were not normalized into a common group space.

#### ROI Definition

Data were analyzed within *a priori* regions of interest (ROIs) derived from a population atlas of large-scale resting-state networks(3). These regions are defined on the Freesurfer average cortical surface mesh. To obtain ROI masks in native functional space, we first reverse-normalized the ROI labels using the spherical registration parameters (36) and then transformed the vertex coordinates into the space of the first run using the inverse of the functional-to-anatomical registration. Voxels intersecting the midpoint between the grey-white and grey-pial boundaries were included in the analyses. This produced ROIs that respected the underlying two-dimensional topology of the cortical surface and minimized the contributions of voxels lying outside of grey matter. To estimate distances between voxels along the cortical surface, we established a one-to-one mapping between voxels and surface vertices. A small number of vertices were originally mapped to multiple voxels; in these cases, we used the voxel whose center was closest to the vertex coordinate. We then constructed a distance matrix between each voxel center where distances were measured with Djikstra’s algorithm on the midthickness cortex mesh (37).

#### Decoding Analyses and Context Preference Estimation

We used linear classifiers to decode information about task context and estimate each voxel’s context preference. Decoding was implemented in three steps. We first used voxelwise models to deconvolve the amplitude of the evoked response in different experimental conditions. The estimated response amplitudes were then used as samples in the multivariate analyses. Finally, we inverted the resulting decoding models to obtain estimates of each voxel’s context preference.

For Experiment 1, the deconvolution model included 12 regressors reflecting a crossing between context (color or motion), trial type (early-cue, concurrent-cue, or cue-only) and cue pattern. For Experiment 2, the deconvolution model included 16 regressors reflecting a crossing between context (color or orientation), cue shape, color stimulus strength, and orientation stimulus strength. These regressors were dummy coded such that for each trial there was only one regressor event. Regressors were defined as boxcars onsetting at the time of the cue and lasting for a duration equal to the sum of the cue duration and mean RT, where applicable. To control for potentially confounding RT differences between contexts (38), we also included parametric regressors where the amplitude of the boxcar was determined by the Z-scored RT on each trial. RT effects were modeled separately for each context and, in Experiment 2, each stimulus strength. These condition regressors were convolved with a canonical difference-of-gammas model of the hemodynamic response function (HRF). We additionally included a set of identical regressors that were convolved with the temporal derivative of the HRF. After assembling and high-pass filtering the design matrix, the confound model that had been used to denoise the data was regressed out. Finally, the deconvolution model was fit to the preprocessed timeseries from each voxel using Ordinary Least Squares, separately for each run, producing a vector of parameter estimates (12 for Experiment 1 and 16 for Experiment 2). The vectors corresponding to each of the voxels from the two ROIs were then stacked, and the vectors corresponding to each voxel were Z-scored across conditions. In total, there were 144 samples for Experiment 1 and 192 samples for Experiment 2.

Multivariate analyses were performed using L2 penalized logistic regression models (39, 40) trained in a binary classification problem to predict task context (motion vs. color trials in Experiment 1 and orientation vs. color trials in Experiment 2). Specifically, this involved minimizing the following cost function:

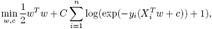

where *y_i_* is the binary label for condition *i*, *X_i_* is the sample vector of estimated BOLD amplitude across the *n* voxels in the ROI, and *C* is a penalty parameter which we set to the default value of 1. We evaluated the accuracy of the decoding models using leave-one-run-out cross-validation and assessed the significance of the results in a permutation framework where we randomly shuffled the context labels, within run, and then re-trained and tested the classifier on each of 100 iterations to establish a distribution of accuracy scores under the null hypothesis (41). For group inferences, we subtracted the mean of each subject’s null distribution from the observed accuracy value and then performed a one-sample *t* test against 0. For subject inferences, we used the percentile of the observed value in the null distribution to obtain a *P* value.

The weights that are estimated when training the decoding model are not directly interpretable, but they can be made interpretable by multiplying the weight vector *w* with the data covariance matrix, Ʃ_*x*_, to produce a vector of context preferences (42):

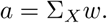

The resulting vector, *a*, is a distributed pattern corresponding to each voxel’s relative preference for the two contexts. To identify voxels with strong context preferences, we compared each voxel’s preference value, *a_j_*, to the vector of preferences assigned to voxel *j* in the permutation test. Voxels where the observed value fell below the 10th percentile or above the 90th percentile in the null distribution were used in subsequent analyses. Note that this procedure uses a liberal threshold as our goal was not to reject a null hypothesis in any individual voxel but rather to identify a suitably large population of voxels that expressed relatively strong preferences for each of the contexts. Our results did not depend on the specific threshold chosen.

To evaluate the spatial organization of the context representations, we determined how well we could predict the preference of a given voxel from the preferences of its neighbors at a given radius. Specifically, we quantified the error at radius *i* as the mean squared difference between the preference in each of the *n* ROI voxels and the mean of the *m* voxels that were separated from it by a distance within 2mm of the specified radius:

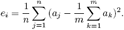

We then compared the observed error value to a null distribution of errors obtained by randomly shuffling the voxel preferences. To quantify the clustering, we estimated an upper bound for the neighborhood size as the first point at which the observed error values were greater than the mean of the null distribution at that distance.

#### Estimation of Spontaneous Correlation Strength

We estimated the magnitude of spontaneous correlations in both task-engaged and resting states. Spontaneous activity in a task-engaged state was estimated using residual data after regressing out a model of task-evoked effects (11, 25). Specifically, the task models were structured identically to the models used during the deconvolution step of the decoding analyses, with the difference that, rather than convolving regressors with a canonical HRF model, we used a finite-impulse-response (FIR) basis to more flexibly fit and remove task-evoked responses. In Experiment 1, where the design was oversampled relative to the data acquisition, we first upsampled the timeseries data to 1 s resolution using cubic spline interpolation. The sampling rate of the design of Experiment 2 was matched to that of the data acquisition and so no resampling was applied. In each case, we used 24 FIR regressors per condition, onsetting 2 TRs before the cue. These models were fit using Ordinary Least Squares to denoised timeseries data that had been concatenated across runs. Analyses of spontaneous correlations in the resting state were performed on the denoised timeseries data. Spontaneous correlation matrices were created for both datasets by estimating the Pearson correlation between the timeseries from each voxel within each run and then averaging correlation values over runs.

We used the data from Experiment 2, where we collected both task-based and resting-state data from the same subjects, to evaluate the similarity of the spontaneous correlation structure under these two conditions. Specifically, we vectorized the upper triangles of the two correlation matrices and then computed a Pearson correlation between the two vectors. We tested the significance of this value with the Mantel Test, where, on each of 100 iterations, we applied the same permutation to the rows and columns of the resting-state correlation matrix and then recomputed the similarity to the task-engaged correlation matrix to generate a null distribution of similarity values.

#### Relating context preferences to spontaneous correlations

To visualize the relationship between context preference and spontaneous correlation, we submitted the spontaneous correlation matrix to metric multidimensional scaling (MDS), which embeds the similarity matrix in a 2 dimensional space. We then created scatterplots where each point corresponded to a voxel. The position of the points was determined by the MDS embedding coordinates, and the color of the points was determined by their context preferences. Because the axes returned by MDS are arbitrary, we computed the matrix-vector product between the MDS coordinates and the voxel preferences and then applied a rotation so that the resulting coordinate fell on the positive side of the *x* axis.

To quantify the relationship between these two variables, we computed the mean correlation between voxels with the same context preference and the mean correlation between voxels with different context preferences. We then evaluated the significance of this value with a permutation test. On each of 100 iterations, we randomly shuffled the context preference labels and recomputed the two measures. We then compared the observed difference in correlations to the distribution of differences under the null hypothesis.

To determine the spatial extent of the functionally selective spontaneous correlations, we recomputed the above measures while systematically excluding correlations between voxels that were separated by less than a specified distance threshold. Note that this is similar to how we evaluated the spatial clustering of the voxel preferences, but here we used all voxels separated by a distance larger than the threshold rather than voxels situated at that specific distance.

#### Code Availability

Data were analyzed using published open-source software and custom code written in Python. Imaging data were processed with a workflow of FSL 5.0.8 (43) and FreeSurfer 5.3.0 (44) tools implemented in Nipype 0.11.0 (45). The Python code used a number of libraries including numpy, scipy, matplotlib, seaborn, pandas, and jupyter. Cortical surface visualizations were created using pysurfer. All custom code will be made available at https://github.com/WagnerLabPapers.

## Supplemental Methods

### Stimuli and Experimental Design: Experiment 2

Subjects in Experiment 2 performed a context-dependent perceptual discrimination task (Supplemental Figure 1B). The stimulus was a 5.5° radius circular array of 0.3° × 0.1° “sticks” centered on a 0.2° fixation point. No sticks were drawn within 1° of fixation. The stick centers were chosen using a poisson disc sampling algorithm to ensure that sticks were densely positioned without overlapping (minimum distance between stick centers: 0.35°) or conforming to a rigid grid. Each stick was oriented 45° left or right of vertical and colored either red or green. Colors were chosen using the CIE I_ch system to obtain distinct hues (red: 60, green: 140) with equal chroma (30) and luminance (~80). The specific luminance values were calibrated before each session with a separate color discrimination task that employed a staircase to match the perceived brightness of the colors. The screen background was drawn in grey at 30% luminance. During the scanning session, the stimulus was presented on a 1920 × 1080 pixel LCD display with a 60 Hz refresh rate, which was placed at the back of the scanner bore and viewed through a mirror attached to the head coil. Behavioral sessions were performed on MacBook Pro laptop computers. Stimulus presentation and response collection were controlled using PsychoPy.

The subject’s task was to make either a color or an orientation discrimination on each trial. On color trials, they had to determine whether more sticks were either drawn in red or green; on orientation trials, they had to determine whether more sticks were oriented left or right. The trial type (the context) was cued with a 3-6 sided 1° wide polygon drawn at 40% luminance around the fixation point. Two distinct shapes cued a color trial, and two shapes cued an orientation trial. These assignments were counterbalanced across participants, and task demands did not otherwise vary between the two cues used for each context. Subjects were instructed to maintain fixation, and eye movements were monitored with a camera inside the scanner bore. Each trial began when the fixation point turned from black to white for 720 ms to signal the imminent trial. The cue and stimulus then appeared simultaneously. Subjects were instructed to respond as soon as they had reached a decision, balancing speed and accuracy Responses were made using button boxes held in the left and right hands. The same buttons were used for both contexts. The mapping between color responses and hands was counterbalanced across participants, but left and right buttons always mapped to left and right orientation decisions. The stimulus disappeared when the subject made a response. Feedback was provided by blinking the fixation point at 10 Hz for 500 ms after errors. Following the trial, the fixation point turned black to signal the inter-trial interval (ITI). The ITI duration was drawn from a truncated geometric distribution over TRs with *p* =  and a maximum of 10 TRs (mean ITI: 1.86 TRs). Each task run had a total duration of 370.8 s.

The stimulus changed dynamically over the course of each trial. This was intended to decrease reliance on a strategy of searching for clusters of sticks with the same feature and, instead, to encourage integration of evidence over time. Specifically, individual sticks were replaced, possibly in a different color or orientation, with a probability of 0.05 on each screen refresh. During replacement, the stick disappeared for 9 frames and then had a 0.5 probability of being redrawn on each subsequent frame. Stimulus strength was manipulated by controlling the proportion of sticks drawn with each color and orientation. We defined stimulus strength as P_feature_ – 0.5; that is, a trial where 60% of the sticks were red had a color strength of 0.1.

Subjects were trained on the task in a pair of 1-hr behavioral sessions that occurred 1-3 days prior to scanning. Subjects first performed the task with high stimulus strength to learn the meanings of the cues and the correct response mappings. This session began with blocked trials that shared a context, but the block length was decreased over the course of the introductory session so that subjects were gradually acclimated to context switching. Subjects then continued the high-stimulus strength practice until they reported comfort with the basic elements of the task and had reached near-perfect accuracy (subjects performed ~650 trials of the practice task). They next performed two blocks of a calibration task (400 trials per block) where the stimulus strength was randomly varied between 0.02 and 0.18 in steps of 0.04 in order to find values to use during scanning. We fit cumulative Weibull functions to the resulting choice accuracy data, separately for color and motion trials, and chose stimulus strengths for hard and easy trials that would be expected to produce 65% and 95% accuracy, respectively

Each task run consisted of 64 trials, representing a full crossing of context (color or orientation), context cue, color strength, orientation strength, dominant color, and dominant orientation. Event schedules for each run were optimized in two steps. First, a large space (n = 50,000) of possible event orders was searched, and the 16 schedules that best minimized the average deviation from the ideal first-order transition matrix within each of the six trial features were selected. Second, “null” events were inserted into each schedule to create a jittered distribution of ITIs that optimized the efficiency of estimating the hemodynamic response function. For each subject, we used 12 of these schedules, in a random order, over the course of the experiment.

**Figure 1:**
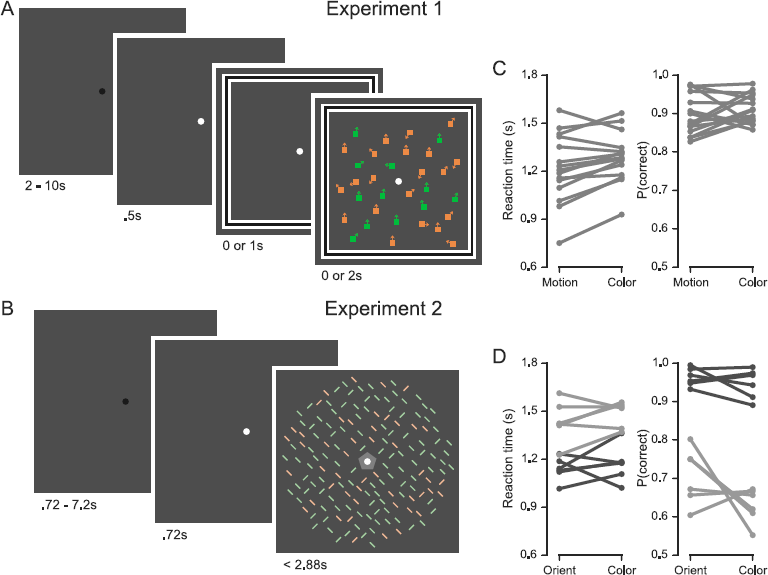
Task design and behavior. (A-B) Experimental designs for Experiments 1 and 2. (C) Mean RT and choice accuracy are shown separately for each subject in Experiment 1. (D) Mean RT and choice accuracy are shown separately for each subject In Experiment 2. Dark and light colors indicate behavior on trials with high and low stimulus strength, respectively.

**Figure 2:**
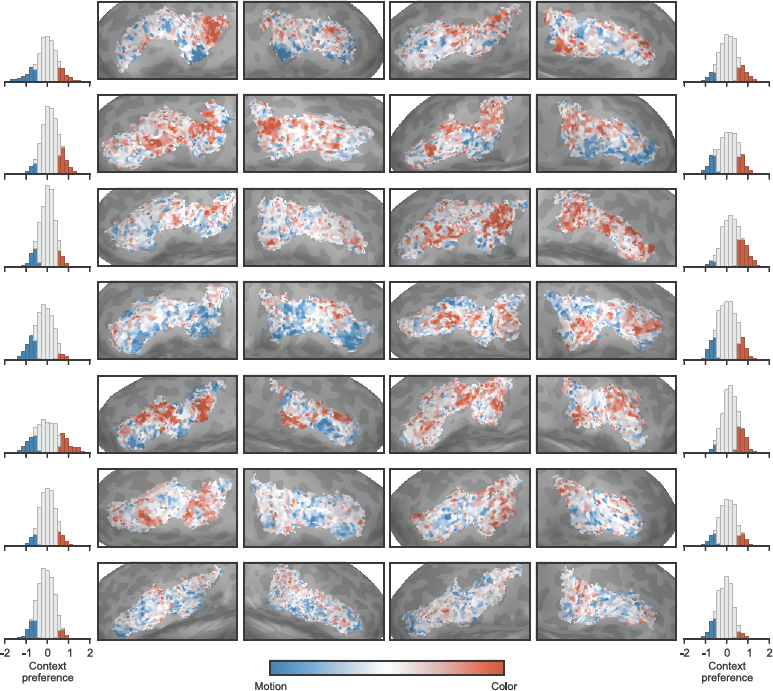
Voxelwise context preferences for all subjects in Experiment 1. Conventions are as in Figure 2. Subjects are ordered (left-to right and top-to-bottom) by decoding accuracy.

**Figure 3:**
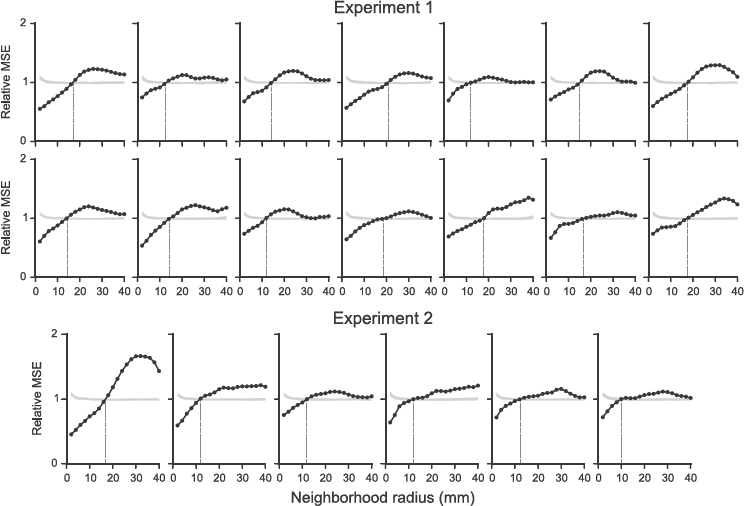
Voxelwise context preferences exhibit relatively fine-scaled spatial interdigitation. Black lines show the mean squared error predicting the preference in ROI voxels by the average of the preferences at the specified radius. Gray bands show the 95% prediction interval from a permutation test where we randomly shuffled voxel preferences. Plots are scaled relative to the mean of the null distribution for ease of visual comparison. The vertical line shows the first point at which the observed error is larger than the error in the null distribution. Significantly lower error before this point provides evidence of spatial clustering, while significantly higher error after this point provides evidence of spatial interdigitation. Each plot shows data from one subject, and subjects are ordered by decoding accuracy.

**Figure 4:**
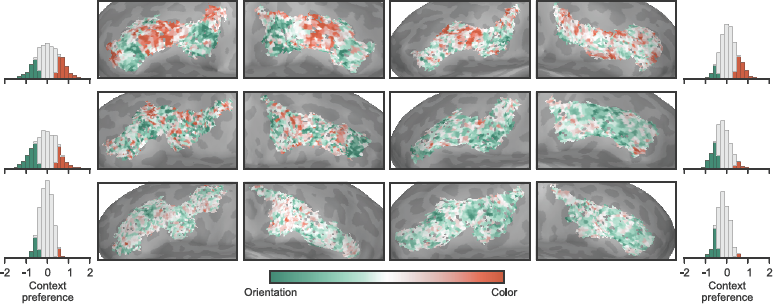
Voxelwise context preferences for all subjects in Experiment 2. Conventions are as in Figure 2. Subjects are ordered (left-to right and top-to-bottom) by decoding accuracy.

**Figure 5:**
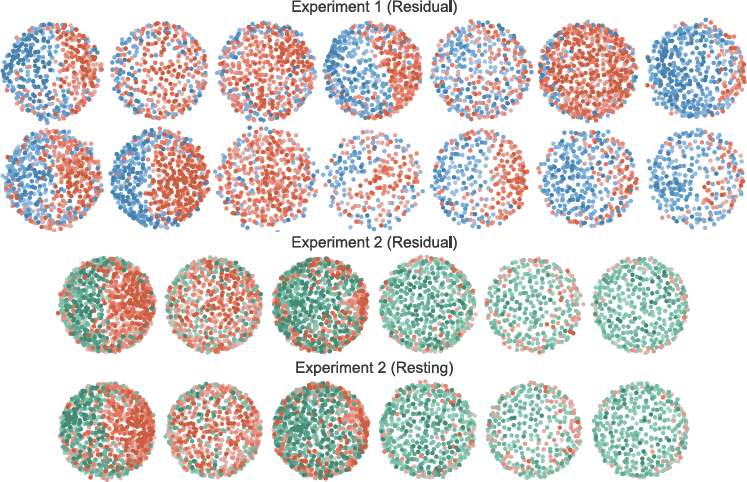
The relationship between spontaneous correlation structure and context preference for all subjects. Conventions are as in Figure 3. Each plot shows data from one subject, and subjects are ordered by decoding accuracy.

**Figure 6:**
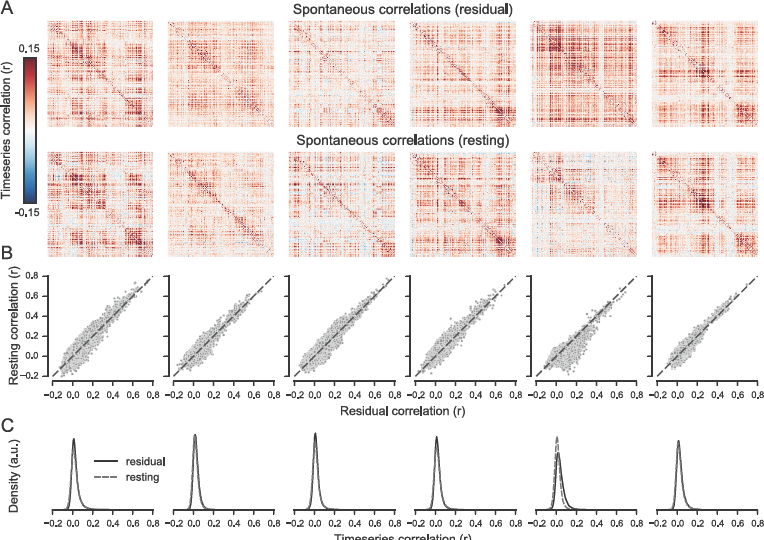
The relationship between spontaneous correlations estimated using residuals in task-engaged state or during rest. (A) Spontaneous correlation matrices for IFS voxels estimated from task-engaged or resting-state data. (B) Scatterplots showing the similarity between the upper triangles of the two matrices in panel A. (C) Density plots showing the distribution of correlations across all pairs of voxels in the IFS. Each column shows data from one subject, and subjects are ordered by decoding accuracy.

**Figure 7:**
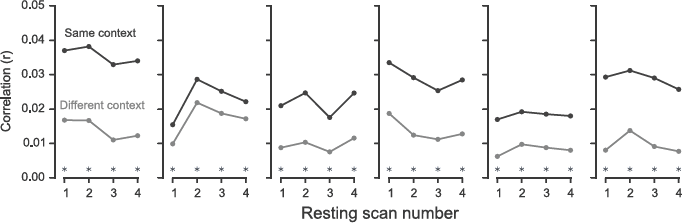
Subnetworks are stable in resting-state scans across the duration of the scanning sessions in Experiment 2. Spontaneous correlations were computed separately for each of the four resting scans in each session and then averaged over sessions. Asterisks indicate significant differences at *P* < 0.05 as determined using a permutation test. Each plot shows data from one subject, and subjects are ordered by decoding accuracy.

**Figure 8:**
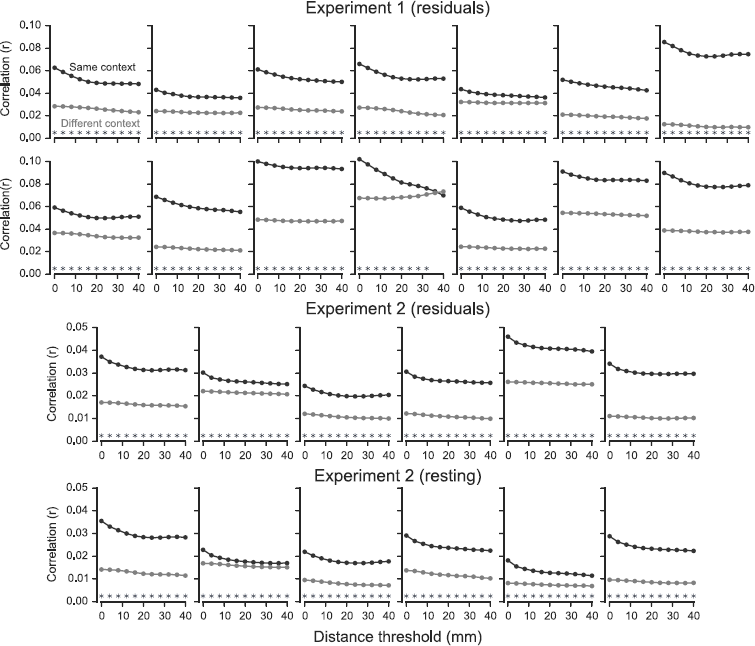
Subnetworks extend over relatively long distances. Lines show the mean correlation between voxels with the same or different context preferences computed while excluding voxels that are more proximal than the specified distance threshold. Asterisks indicate significant differences at *P* < 0.05 as determined using a permutation test. Each plot shows data from one subject, and subjects are ordered by decoding accuracy.

